# High-Content, Cell-by-Cell Assessment of HER2 Overexpression and Amplification for Heterogeneity Detection in Breast Cancer

**DOI:** 10.1101/419499

**Authors:** Huu Tuan Nguyen, Daniel Migliozzi, Bettina Bisig, Laurence de Leval, Martin A.M. Gijs

## Abstract

Immunohistochemistry and fluorescence in situ hybridization are the two standard methods for Human Epidermal Growth Factor Receptor 2 (HER2) assessment. However, they have severe limitations to assess quantitatively intratumoral heterogeneity (ITH) when multiple subclones of tumor cells co-exist. We develop here a high-content, quantitative analysis of breast cancer tissues based on microfluidic experimentation and image processing, to characterize both HER2 protein overexpression and *HER2* gene amplification at the cell level. The technique consists of performing sequential steps on the same tissue slide: an immunofluorescence (IF) assay using a microfluidic protocol, an elution step for removing the IF staining agents, a standard FISH staining protocol, followed by automated quantitative cell-by-cell image processing. Moreover, ITH is accurately detected in both cluster and mosaic form using an analysis of spatial association and a mathematical model that allows discriminating true heterogeneity from artifacts due to the use of thin tissue sections. This study paves the way to evaluate ITH with high accuracy and content while requiring standard staining methods.

## Introduction

In 15-20% of breast cancer cases, the human epidermal growth factor receptor 2 (HER2, also known as ERBB2 or HER2/neu) is overexpressed, causing rapid progression and poor prognosis of the disease (1). This cancer subgroup (HER2 positive) favorably responds to HER2-targeted therapies (e.g., trastuzumab, pertuzumab, lapatinib and trastuzumab emtansine). According to the American Society of Clinical Oncology (ASCO) / College of American Pathologists (CAP) guideline recommendation in 2013 (1), IHC and FISH are two validated techniques for HER2 assessment. Conventional IHC is inherently subjective and qualitative, as the evaluation relies on the experience and judgment of the pathologist. Interpretation difficulty in IHC can be a source of diagnostic errors (1, 2). Compared to IHC, FISH is more quantitative, but only a small tumor area, corresponding to 20–100 cells, is usually manually scored to evaluate the HER2 status. More importantly, assessment of HER2 intratumoral heterogeneity (ITH) is challenging for both methods, as it is characterized by differences in HER2 status among different subclones and cells in different regions of a tumor (3). HER2 ITH is often associated with poor prognosis and resistance to HER2-targeted therapy (4). Two forms of HER2 ITH exist: coexistence of discrete focal clones of cells (*i.e.*, “cluster” heterogeneity) or individual cells placed in a dominant background of the opposite status (*i.e.*, “mosaic” heterogeneity) (5). We consider a mosaic heterogeneity as an admixture of a positive clone within a negative cluster or a negative clone in a positive cluster (6). In clinical assessment, the presence of “cluster” ITH can be confirmed, if the proportion of the area of a minority cellular phenotype within that of the majority is above 10% (1). The HER2 “mosaic” heterogeneity is confirmed if the proportion of heterogeneous cells, as obtained by the manual scoring of *in situ* hybridization (ISH) signals, is between 5-50% of all cancer cells scored (6). In case of a small number of cell counts, the statistical power of the obtained percentages of heterogeneous cells is low.

To improve HER2 overexpression assessment, some researchers used automatic IHC scoring software (7). Some other studies have proposed using multichannel computational analysis of IF images to quantify the HER2 protein presence (8-10). Our group has demonstrated that combining microfluidics and digital IF quantification can improve diagnostic accuracy (9, 10). For FISH analysis, automatic counting was developed to decrease the image analysis time and reduce human errors during FISH scoring (11-17). However, the used high-magnification objectives (63×) with a small depth of focus require taking z-stack images for different focal planes for recording FISH signals, so that the necessary computational power and memory requirements during image processing are large, still resulting in a small area that could be analyzed.

Here, we develop a new method based on microfluidics and image processing for high-content combined analysis of HER2 protein overexpression and *HER2* gene amplification in large breast cancer tissue areas. In each cell, we quantify the HER2 protein intensity and its background, CK protein intensity (obtained from IF), the number of *HER2* gene loci and CEP17 (obtained from FISH), and cell positions. The whole slide is recorded and analyzed by a low magnification (20×) objective and image processing software, allowing automated evaluation of 10^4^−10^5^ cells. We demonstrate that both cluster and mosaic ITH can be detected and quantified in a large tissue area based on the local indication of spatial association (LISA) method (18). This technique, widely used for spatial analysis of geographical datasets, is a powerful statistic tool explicitly adapted to our specific problem of cell-to-cell variability in a tissue. Finally, we achieve a quantitative estimation of cluster and mosaic ITH in a small cohort of 20 clinical breast cancer tissue slides. Using a numerical model, we could discriminate true mosaic ITH from variations of *HER2* loci in a cell cut as caused by truncation artifacts.

## Results

### Cell-by-cell characterization using microfluidic immunofluorescence and automated FISH scoring

The protocol for microfluidic HER2 and CK IF staining was described previously (9, 10). Briefly, after standard removal of paraffin and antigen retrieval, the immunostaining of a tissue slide was performed using a microfluidic chip flushed respectively primary Ab probes, washing buffer and detection secondary Abs on the tissue slide surface (Fig. 1ai, ii and section S1 of the SI). After the IF staining step and 4’,6-diamidino-2-phenylindole (DAPI) mounting medium application, the whole surface of the slide was successively imaged and treated with a proteolytic enzyme to remove all proteins before performing HER2 DNA FISH (Fig. 1aiii-iv and section S2 of the SI). In the following step, cells in the IF image were then segmented using image processing software (Fig. 1bi). The HER2 intensity was then enhanced and thresholded, defining the membrane area where HER2 and CK signals were measured. In the FISH images, the regions of interest defining the nuclei were determined (Fig. 1bii) and associated with cells identified in the IF image. IF and FISH signal characteristics, such as immunostaining intensity and FISH scores, were obtained and merged into a single database (Fig. 1c). For tissue analysis, the data processing was followed by a spatial analysis (*vide infra*). All tissues in the batch are successfully stained and analyzed.

**Fig. 1:**
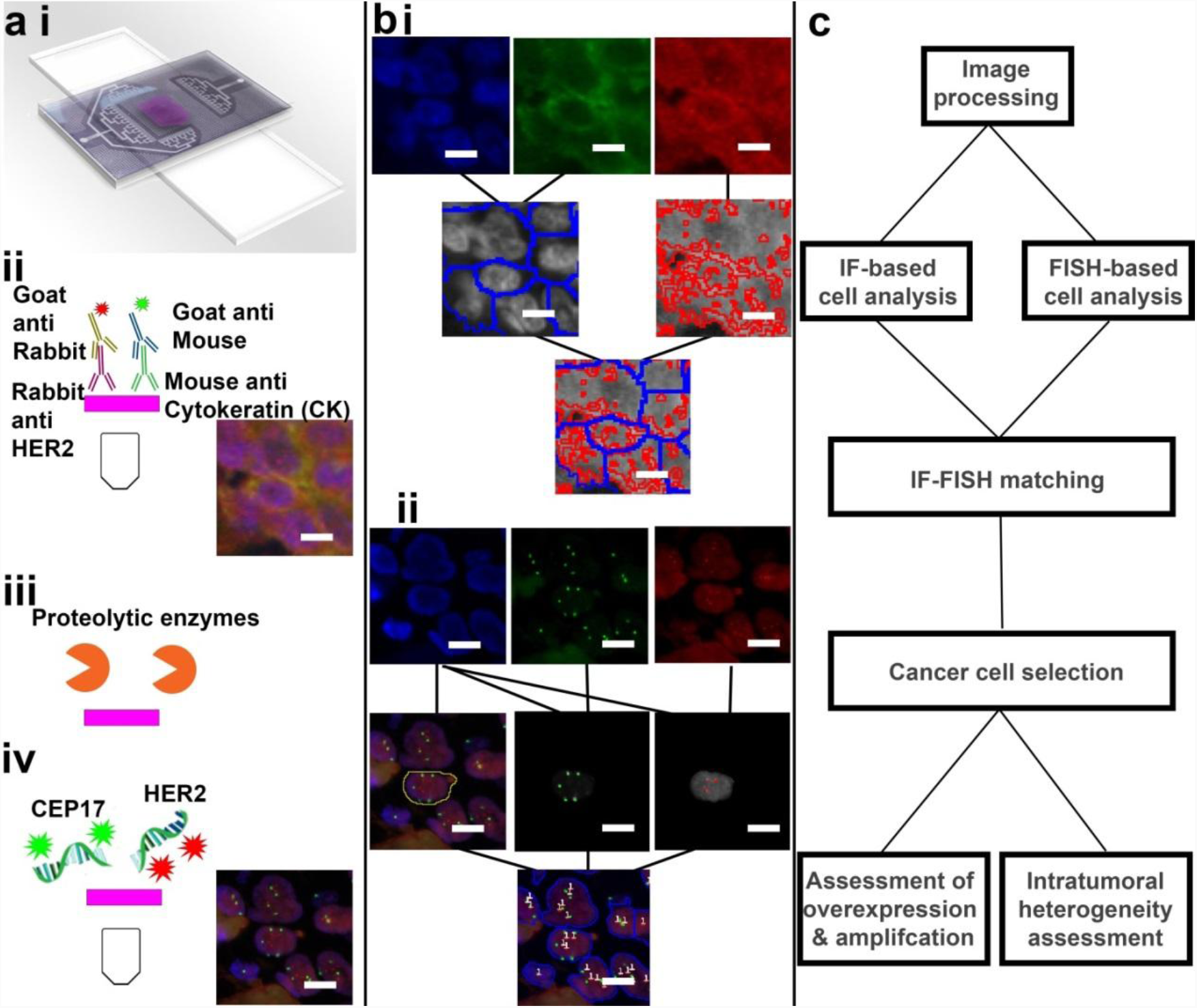
High-content analysis of microfluidic immunofluorescence (IF) and fluorescence *in situ* hybridization (FISH). **(a)** Sequential IF/FISH staining. (**i)** Immunostaining of cell pellet or tissue section via the microfluidic tissue processor (MTP) clamped onto the tissue-carrying glass slide to deliver the reagents in a highly-controlled fashion. (**ii)** In the IF protocol, human epidermal growth factor receptor 2 (HER2) and cytokeratin (CK) proteins are tagged with rabbit anti-human HER2 antibody and mouse anti-human CK antibody and detected using AF594-labeled goat anti-rabbit IgG antibody (Ab) and AF647-labeled goat anti-mouse IgG Ab, respectively. Nuclei are marked with DAPI. The whole slide is scanned using a low magnification objective (20×). The image of a cluster of cells is presented (HER2: red, CK: green, DAPI: blue). **(iii)** In an elution step, staining agents are removed from the slide using proteolytic enzymes. **iv** In the FISH protocol, the *HER2* loci are labeled with fluorescent *HER2* probes, the centromeres of chromosome 17 are labeled with fluorescent chromosome enumeration probes (CEP17), and the nuclei are marked with DAPI. An image of the same cells as in ii is shown (*HER2* loci: red, CEP17: green, DAPI: blue). **(b)** Image-processing of the IF and FISH images obtained after the protocol from **A**. IF and FISH images, aligned using the common DAPI channel, are sequentially processed. (**i)** For IF analysis, clusters of cells are segmented into an individual cell or a smaller group of cells based on nuclear analysis from the DAPI and CK channels. The HER2 signal also defines the area in which the mean HER2 and CK intensities for each cell are measured. (**ii)** For FISH analysis, nuclei define the area where HER2 and CEP17 signals for each cell are scored. **(c)** Data analysis pipeline. The cell HER2 expression given by IF is merged with the cell HER2 amplification obtained from FISH, followed by a filtering step that selects the cells of interest. For each sample, scores for HER2 protein overexpression and *HER2* gene amplification based on the analysis of all the cells are obtained. Intra-tumoral heterogeneity analysis using spatial association analysis is also performed. Scale bars: 10 µm.

### High-content analysis of HER2 overexpression and amplification in cell lines and tissues

#### Cell line overexpression and amplification analysis

To assess the robustness of HER2 overexpression and amplification with our technique, we perform IF/FISH staining and analyses on a HER2-positive cell line (SKBR3) and a HER2-negative cell line (MD-MB-468). Mean values of HER2 intensity from an IF image and automatic *HER2* loci per cell (*HER2* loci/cell) scores from FISH are plotted in Fig. S1 of the SI. While a wide range of variation of HER2 IF signals among cells is observed, the means HER2/CK ratio of all cells are similar between the replicates. Also, less variation and a sharper distinction between negative and positive cell lines are observed (see Fig. 2a), suggesting that the HER2/CK ratio can be used to characterize each cell line sample. For FISH, we also observe a large *HER2* loci/cell variation among cells of a slide (Fig. S1), which is explained by truncation artifacts as cells were sliced into thin sections. Thus, we normalize the *HER2* copy number to the CEP17 number for each cell and calculate the average *HER2*/CEP17 ratio of the whole cell population to cancel the truncation effects. Finally, the two selected parameters plotted along the axes of Fig. 2a are the mean HER2/CK ratio and the mean *HER2*/CEP17 ratio. In a first approach, we propose the threshold between HER2-negative/equivocal and positive tissues for quantitative IF as the lower bound of at-test’s 95% confidence interval of the mean HER2/CK ratio of the SKBR3 cell line. This gives a threshold 0.25 for IF HER2 positivity classification in tissues (*vide infra*).

**Fig. 2:**
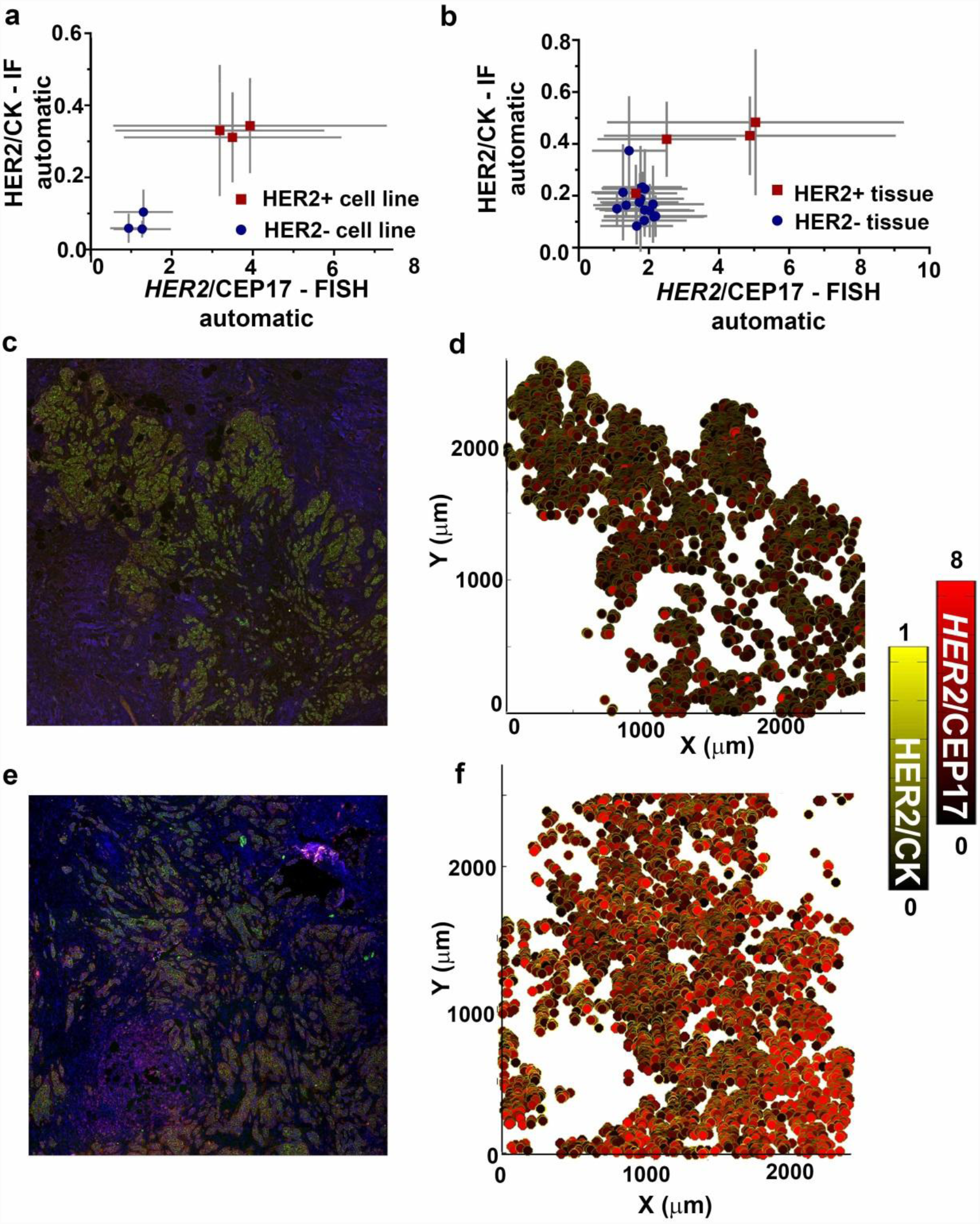
HER2 assessment in cell lines and tissues by using automatic and quantitative IF-FISH analysis. **(a)** HER2 overexpression (given by the cell-by-cell ratio between the HER2 and the CK signals) versus HER2 amplification (given by the cell-by-cell ratio between the number of *HER2* loci and CEP17) for HER2 positive (+) and HER2 negative (-) cell lines. Data are plotted as a mean ± standard deviation. **(b)** Assessment of HER2 overexpression and amplification in HER2+ and HER2-tissues with the same methodology as in **A**. The HER2 status is obtained from assessment by a pathologist using a standard FISH technique. The error bars represent the variation of these scores among cells in the tissue. **(c)** IF image of a HER2-equivocal tissue. **(d)** Cell-by-cell representation of HER2/CK. and *HER2*/CEP17 signals. Cells are represented by dots. The contours of the dots (in yellow scale) denote the normalized HER2/CK ratio, while the inside of the dots (in red scale) indicate the *HER2*/CEP17 ratio of the cells in **C**. Normalization of the HER2/CK ratio (yellow scale) is obtained by allocating the value 1 to the maximum HER2/CK ratio that was obtained from all tissues. The maximum of the red scale is chosen as 8 for easy distinction between positive and negative cells. **(e)** IF image of a HER2-positive tissue. **(f)** Cell-by-cell representation of the tissue in **E** with the same methodology as in **D**. Colors in IF images: Blue=DAPI, Green=CK, Red=HER2.

#### Tissue analysis using the high-content IF-FISH technique

Combining the IF and FISH automatic analyses, all mean HER2/CK ratios obtained from IF and *HER2*/CEP17 ratios obtained from FISH were computed for a small cohort of breast cancer cases, see Table S1 of the SI. The results are plotted in Fig. 2b. In general, we observe a higher range of HER2/CK ratios and *HER2*/CEP17 ratio in the HER2-positive cases compared to that in the equivocal/negative cases, except for one positive case (tissue 1, Table S1) which has significantly lower HER2/CK ratio and *HER2*/CEP17 ratio and one equivocal case (tissue 17, Table S1) which has significantly higher HER2/CK ratios. The positive case is a cluster heterogeneity case. It has a score that varies strongly depending on the location and size of the region of interest where it is assessed. The equivocal case is a metastatic case of breast cancer in bones, for which CK staining is weaker than usually, which increases the HER2/CK ratio. A cell-by-cell representation of a HER2-equivocal tissue is shown in Fig. 2c. For the HER2-equivocal tissue (Fig. 2c), both the HER2/CK ratios are codified by the yellow scale (contours, Fig. 2d), and the *HER2*/CEP17 ratios are codified by the red scale (inside of the dots, Fig. 2d). These parameters are lower than these same parameters for a HER2-positive tissue (Fig. 2E,F).

### Accurate HER2 status assessment based on quantitative IF/FISH

To evaluate its performance compared to the conventional diagnostic methods, we studied the correlation between 3 scores obtained by our method (i.e., HER2/CK ratio, *HER2* loci/cell number, and cell *HER2*/CEP17 ratio) and the pathologist’s scores (i.e., standard *HER2* loci/cell number and overall *HER2*/CEP17 ratios). We benchmarked our method with *HER2* loci/cell and *HER2*/CEP17 ratios instead of the IHC scores because FISH is more quantitative than IHC. In Fig. 3a, the HER2/CK ratio obtained by automatic IF analysis was compared to the standard *HER2*/CEP17 scores obtained from manual counting. Here, we obtain a sum-of-square R^2^ equal to 0.64. The dotted lines are the thresholds, defined as follows. For IF, we obtained from cell line analysis a threshold of 0.25 for positivity determination. For the overall *HER2*/CEP17, we used the same threshold as in standard diagnostics routine (see section S3 of the SI), which is equal to 2. By using these thresholds, all but two cases are correctly classified. These outliners are mentioned previously (cases of bone metastases and heterogeneity). Without these two particular cases, R^2^=0.80, suggesting an acceptable correlation between the two scores. In Fig. 3b, the mean number of *HER2* loci/cell obtained from an automatic FISH method is plotted against these obtained via manual scoring (linear regression R^2^= 0.88). If the heterogeneous case is not included in the dataset, R^2^= 0.93, which shows a high correlation between automatic and standard scoring methods for tissue analysis. Finally, in Fig. 3c, the cell *HER2*/CEP17 ratio by automatic FISH also correlates well with the overall *HER2*/CEP17 ratio obtained by the standard FISH method (R^2^=0.81). If the heterogeneous case is not included, R^2^=0.93.

**Fig. 3:**
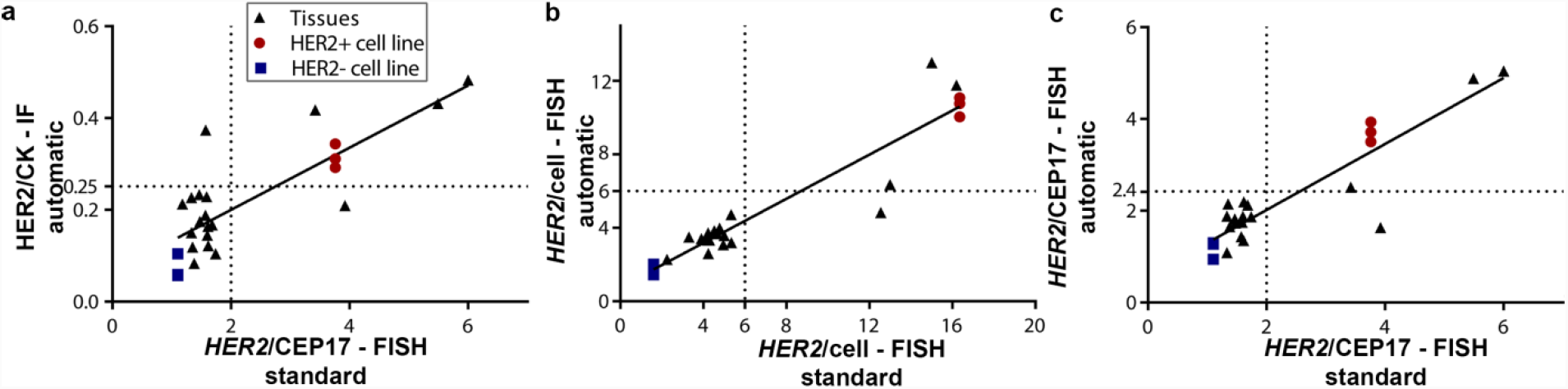
Comparison of quantitative and automatic IF-FISH scoring method with the standard HER2 assessment for tissue and cell line samples. **(a)** Correlation between HER2 overexpression (HER2/CK ratio obtained from the microfluidic staining protocol and automated IF image-processing) and HER2 amplification (*HER2*/CEP17 ratio obtained from standard FISH scoring). The threshold 0.25 for positivity of HER2/CK obtained by IF is defined as the lower bound of the 95% confidence interval obtained from a t-test on the mean HER2/CK scores of three HER2-positive cell line samples. The threshold 2 for positivity of *HER2*/CEP17 obtained by FISH is obtained from the ASCO/CAP 2013 guidelines. **(b)** Correlation between the *HER2* loci number per cell obtained from our automated counting algorithm and the standard FISH technique. The threshold for positivity for the variable *HER2* loci/cell is taken as 6 (dotted lines), obtained from the ASCO/CAP 2013 guidelines. **(c)** Correlation between the *HER2*/CEP17 ratio obtained from our automated counting algorithm and the standard FISH technique. In our automatic method, the *HER2*/CEP17 ratio is calculated as the mean of the *HER2*/CEP17 ratios in all CK-positive cells of a tissue, while in the standard method it is calculated as the ratio of mean *HER2* and CEP17 signals in one or several clusters of 20 cells chosen by the pathologist. The threshold for positivity for the automatic *HER2*/CEP17 ratio (horizontal dotted line) is 2.4 (see text).

### Spatial analysis and evaluation of ITH

#### Cluster heterogeneity

Now we demonstrate that the cluster ITH in the particular heterogeneous case (tissue 1, Table S1) can be effectively identified and quantitatively characterized using the LISA analysis. The LISA method classifies cells based on their own HER2 status (High, Low or non-classified) and the status of their neighbors (High or Low), resulting in: High-High (HH), High-Low (HL), Low-High (LH), Low-Low (LL) and non-classified (NC)-type cells. First, we observe that the IF image of the CK-positive cells in a large area of the tissue displays fairly homogeneous staining of HER2 in red and CK in green (Fig. 4a). The co-presentation of IF overexpression using the HER2/CK ratio (contours in yellow scale) and FISH amplification using the *HER2*/CEP17 ratio (inside of the dots in red scale) of CK-positive cells in the tissue does not display remarkable heterogeneity, as shown in Fig. 4b. However, this is a case having amplification cluster ITH, as proved by LISA analysis of the variable *HER2* loci/cell obtained from the FISH analysis (*vide infra*). In Fig. 4ci, the IF image of a FISH-positive cluster of the tissue is displayed. LISA analysis of the parameter HER2/CK from the IF image shows only LL cells are in this region, confirming that the HER2 expression is clustered and homogeneously negative in this area (Fig. 4cii). However, in the FISH image (Fig. 4ciii) this area is displayed with a majority of HER2-amplified cells, as highlighted by the automatic analysis of FISH signal (Fig. 4civ). In Fig. 4cv, the LISA map for FISH indicates that most cells are HH, meaning that they are amplified (*HER2* loci/cell number>6) and clustered, while two other cells are non-classified (4≤*HER2* loci/cell number≤6). In Fig. 4di, an IF image of a FISH-equivocal cluster (*HER2* loci/cell in between 4 and 6) of the tissue is displayed. While the cells’ IF signal is still classified as negative and clustered (Fig. 4di, ii), the FISH image (Fig. 4diii) shows that most cells are equivocal, except one cell that is highly amplified. The automatic scoring of FISH signal and LISA analysis both confirm that this area has a majority of FISH-equivocal cells and one heterogeneous HL cell (Fig. 4div, v). In Fig. 4E, a LISA map of the IF HER2/CK ratios of CK-positive cells in the tissue area is presented, reconfirming that the staining is fairly homogeneous. In Fig. 4f, LISA maps of the *HER2* loci/cell from the FISH image of the tissue area are plotted. We observe the presence of a FISH-positive subclone (red) and a FISH-negative/equivocal subclone (green). Finally, the proportion of HH, HL, LH and LL cells among all CK-positive cells are obtained for heterogeneity quantification (see table S1). The percentage of HH cells in this HER2-negative-majority tissue is 17% (see tissue 1, table S1), which is higher than the threshold 10%, we can conclude that the tissue has cluster heterogeneity. As another application of the method, we could confirm the positivity and negativity of different cancer components (HER2-positive ductal carcinoma in situ *versus* HER2-negative invasive ductal carcinoma) using the IF-FISH high-content analysis (Fig. S2 in the SI).

**Fig. 4:**
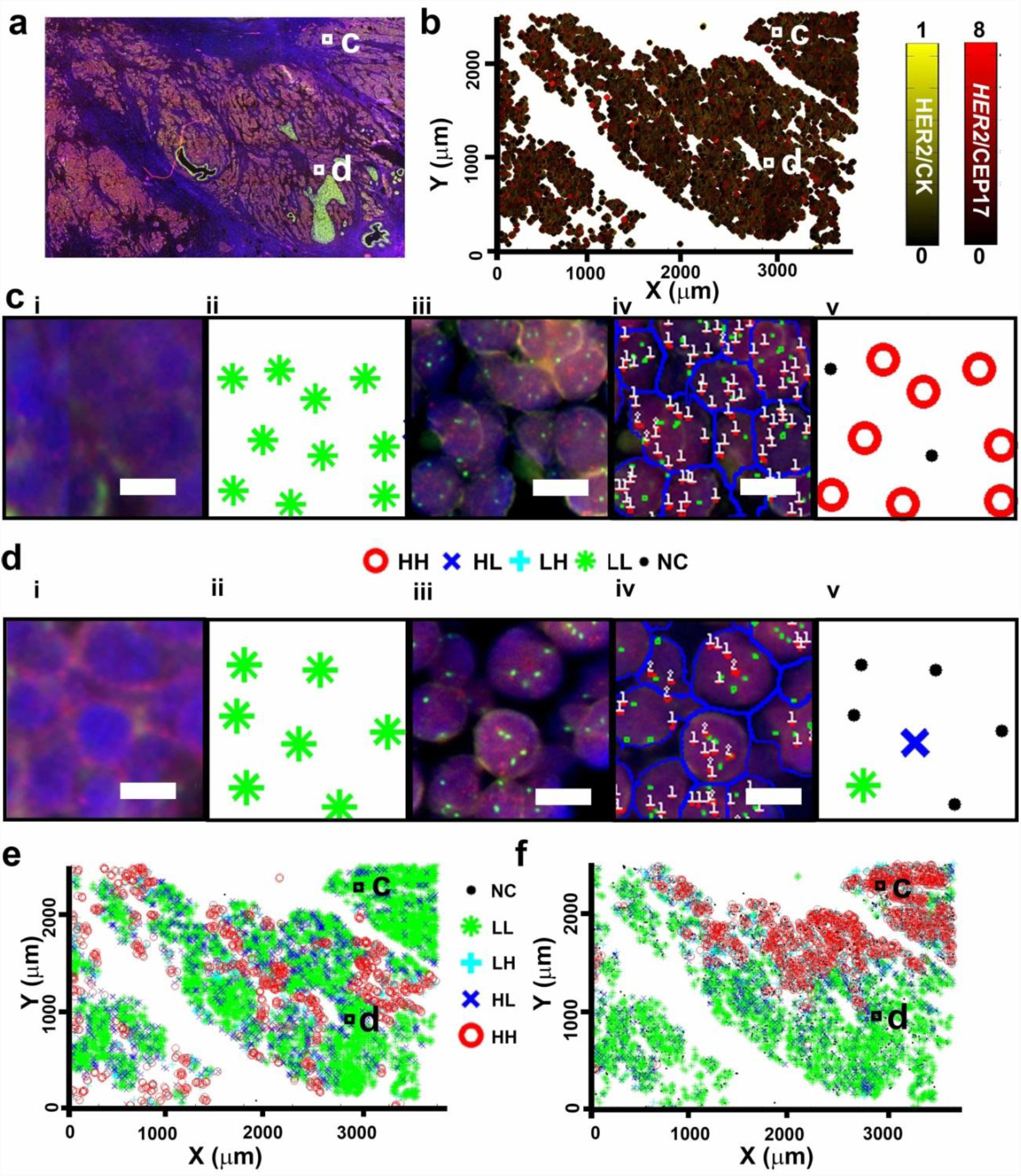
Analysis of a heterogeneous tissue with local indication of spatial association (LISA). **(a)** IF image of a tissue with the definition of two regions of interest C and D, in which we will analyze the heterogeneity in gene amplification. **(b)** Model of the tissue in A using the same cell-by-cell representation as in Fig. 2d,e. **(c)** Heterogeneity analysis of the region of interest c of the tissue in A **i** IF image: blue=nuclei, red=HER2, green=CK. **ii** LISA analysis of the cells in **i**. Cells are classified based on their own HER2 IF-status (High or Low) and on the IF-status of their neighbors (High or Low), resulting in: High-High (HH), High-Low (HL), Low-High (LH), Low-Low (LL)-type cells. Cells and their neighbors are classified as High (respectively Low) if their HER2/CK ratio is higher (respectively lower) than a threshold of 0.25 as obtained from Fig. 3a. **iii** FISH image of the region. Blue=nuclei, green=CEP17, red=HER2. **iv** Automatic scoring of *HER2* loci and CEP17 of the region in **iii**. **v** Spatial association status of the cells in **iv**. Cells are classified as High (respectively Low) if their *HER2* loci number is ≥6 (respectively <4). They are non-classified (NC) in the intermediate interval of *HER2* loci number from 4 to 6, where the FISH HER2 status is equivocal. (**di-v**) Heterogeneity analysis of a HER2 equivocal region of the region of interest d of the tissue in A, following the same procedure as in C. While the IF readout is similar, the FISH status is clearly distinct. **(e)** Spatial association analysis of the HER2 protein expression for the whole tissue in A. **(f)** Spatial association analysis of *HER2* amplification for the whole tissue in A, showing cluster heterogeneity, i.e., having clusters of HH cells that span more than 10% of the tissue area. Scale bars: 10 µm.

#### Mosaic heterogeneity

Following LISA analysis, mosaic ITH is the proportion of HL cells (for a HER2-negative patient) or LH cells (for a HER2-positive patient) in all CK-positive cells. The measured mosaic heterogeneity is then compared to the theoretical model representing the variation of the number of *HER2* loci per cell caused by truncation artifacts in a homogeneous cell population. In Fig. 5a, we present both the truncation artifact model curves (lines) and the measured mosaic ITH in all tissues and cell lines (symbols). By using a statistical model, we demonstrate that even a homogeneous cell population can display apparently heterogeneous cells due to truncation errors. The theoretical curve (figure 5a, lines) representing the truncation artifact in a homogeneous truncated cell population is explained as follows. First, there is no highly amplified cell for the low mean *HER2* loci/cell value. When the number of *HER2* loci/cell increases, the *HER2* loci population shifts to a higher value, thus more truncated cells are recognized as positive. After the mean passes 6, the criterion for selecting heterogeneous cells changes to LH cells (6), resulting in a sharp reduction of the proportion of heterogeneous cells.

**Fig. 5:**
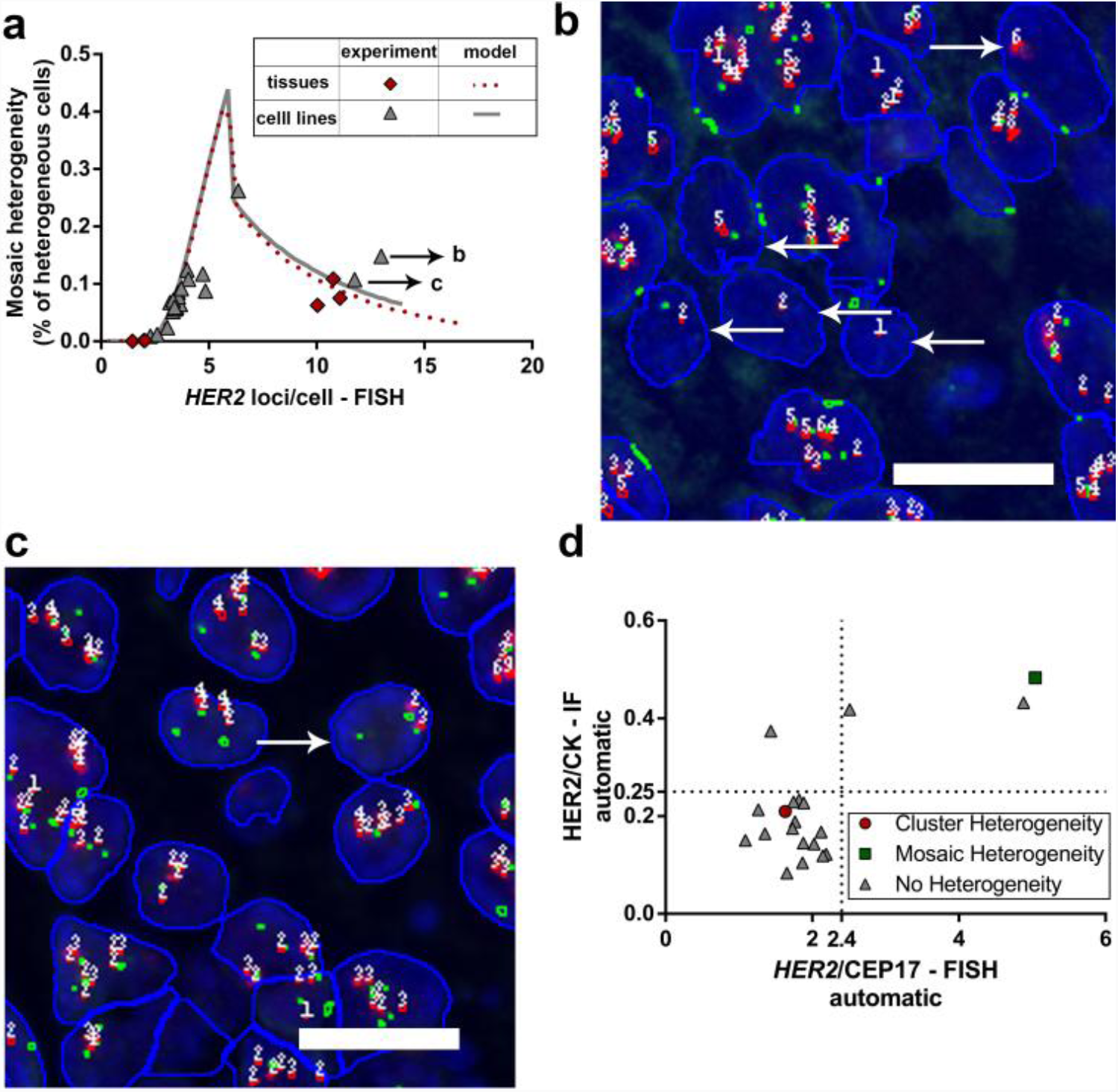
Interpretation of mosaic intratumoral heterogeneity (ITH) by comparison to a statistical model of truncation artifact. **(a)** The mosaic ITH is measured in all tissue samples and benchmarked with the readout of the two cell lines: it is defined as the ratio of the number of individual heterogeneous cells (i.e., cells with positive/negative FISH status in a cluster of cells of negative/positive FISH status), divided by the total number of cancer cells in a tissue. For a HER2 positive sample (mean *HER2* loci/cell≥6 or *HER2*/CEP17≥2), mosaic ITH cells correspond specifically to LH cells of spatial association analysis, like the ones marked by the + symbol in Fig. 4F. For a HER2-negative and equivocal sample (mean *HER2* loci/cell<6 and *HER2*/CEP17<2), mosaic ITH cells correspond to HL cells of spatial association analysis, like the ones marked by the × symbol in Fig. 4F. For both cell lines and tissues, the apparent mosaic ITH of a homogeneous population can be artificial when it is due to truncation artifacts. Indeed, truncated cells having a random position of the nucleus with respect to the cut contain a number of *HER2* dots that follows a binomial law of probability. The full line represents a model simulation that describes such truncation artifacts in a homogeneous population of cells as a function of varying *HER2* loci number. Data points lying above the model curve can be considered as exhibiting true mosaic ITH (see an image of B), while points on the curve represent a false mosaic ITH, as due to truncation artifacts (see an image of C). **(b)** The automatic scoring FISH image of a true mosaic ITH. Arrows indicate cells that differ from their neighborhood. **(c)** The automatic scoring FISH image of an artificial mosaic ITH case. Less heterogeneous cells are observed. Scale bars: 20 µm. **(d)** HER2 overexpression versus *HER2* amplification for all tissues with additional heterogeneity information. It shows one patient with true mosaic ITH (the same as shown in D) and one patient with cluster heterogeneity (data of Fig. 4F) among the 20 patients tested. Thresholds for positivity are indicated by the dotted lines.

The mosaic ITH fits well to the truncation artifact model for most cases. Moreover, when the cell line’s homogeneous population was analyzed by LISA, we find that positive cell lines (Table S1) also have a strong mosaic ITH (from 6 to 11%, the variation can be explained by a smaller number of cells analyzed compared to that of tissues), which confirms the truncation artifact hypothesis. Therefore, if the apparent mosaic ITH is lower or similar than the model, we can thus associate the measured mosaic ITH to the truncation artifacts. Lower mosaic ITH compared to the model is observed in two tissues having the highest percentage of clustered heterogeneous cells (tissue 1 and 13, Table S1). In these two tissues, the mosaic-heterogeneous cells were only considered in the part of the tissues where the majority cell type (HER2-negative) is placed, hence decreasing the overall percentage of the mosaic heterogeneous cells in the whole cell population. Finally, one tissue sample with significantly higher mosaic ITH than the model is potentially exhibiting true mosaic ITH (tissue 19, Table S1). Indeed, we observe in the FISH scoring image that the presence of heterogeneous cells in the true mosaic ITH tissue (Fig. 5b) is more prominent than those in a homogeneous tissue having a similar *HER2* loci/cell score (Fig. 5c). Our method has successfully confirmed the presence of ITH in both cluster and mosaic form. Finally, a simple graph recapitulating the HER2 phenotypic and genotypic characteristics of each tissue as well as their cluster and mosaic heterogeneity can be plotted (Fig. 5d).

## Discussion

In this work, we propose an experimental and analytical method for quantitative assessment of both *HER2* gene amplification and HER2 protein overexpression in a cell-by-cell basis of the same breast cancer tissue slide. The novelty of our method is in the unique combination of (biochemical/computational) steps used for protein and gene biomarker quantification, with application to diagnostics and heterogeneity assessment. First, we demonstrate that the adjusted HER2/CK ratio averaged over segmented cells has a linear correlation with the standard *HER2*/CEP17 ratio. Previous reports also benchmarked to the standard *HER2*/CEP17 ratio and showed that the microfluidic system render a better quantification of the IF signals by keeping the IF signal away from saturation by using short-time flushing of Abs, favoring the discrimination of HER2-positive and -negative cases (9, 10). However, the histogram-based IF scoring method (9, 10) was not standardized because they were not stand-alone variables, but needed to be normalized to a reference signal, for example, a known positive tissue slide (9). Instead, this new IF quantification method does not require the use of benchmark tissues with a known status for staining comparison and only need a low-resolution image. In the future, comparative studies between our IF and other automatic IHC scoring software using the same batch of tissues processed by microfluidic staining can be performed (7). Regarding FISH quantification, our imaging technique uses a 20× objective and an image processing software to analyze the image. This avoids extensive z-stack imaging that is necessary when using a high-resolution 63× objective and allows acquiring signals over a large region of interest of a tissue or cell sample (4 mm × 4 mm in our case). More importantly, this research paves the way to study spatial ITH quantitatively. We highlight that the mechanism responsible for the presence of ITH and its biological consequences are not well understood (20). The main challenge is the lack of a suitable analysis tool that can record and analyze a large number of cells. To address this need, quantitative assessment of ITH has been recently developed, principally based on imaging of IHC and bright-field ISH staining (21-23). However, limitations of the number of markers used, the difficulty of cell segmentation, and non-linearity of IHC signal impede cell-by-cell analysis of both protein expression and gene amplification. Another challenge to address HER2 ITH detection is that most FISH-based clinical studies used the *HER2*/CEP17 ratio for both tissue and cell classification. Although the parameter *HER2*/CEP17 is a good indicator of *HER2* amplification for a population of cells, as it cancels the effect of nucleus truncation, tissue thickness, the mitotic index of the tissue, and abnormal chromosome copy number in some rare aneusomy cases (5), it can give a wrong classification at the cell level due to truncation artifacts of the CEP17 signal. Therefore, in our study, *HER2* loci/cell are used for examination of *HER2* gene heterogeneity (5). Using high-resolution analysis of IF/FISH images, both overexpression and amplification of cells at the same location over a large area of a slide are examined, which increases the accuracy of cluster heterogeneity detection (5). This method is in line with the current recommendation for HER2 ITH testing (5), while being more robust and informative. For mosaic heterogeneity, the preliminary data show that most mosaic heterogeneity isan artifact due to ISH preparation (5, 25). Interestingly, one true mosaic heterogeneous case (tissue 19, Table S1) is successfully identified among all tissues tested, as it has a higher heterogeneous cell proportion than the one corresponding to the truncation artifact model. We believe that mosaic and cluster heterogeneity detection will shed light on the cancer evolution whose cause is still under debate. Two models are widely accepted: cancer stem cell and clonal evolution (3). Hypothetically, the mosaic heterogeneity could be explained by a cancer stem cell model where the two cancer cell types (HER2-positive and HER2-negative) are differentiated from the same progenitor or stem cell population. Therefore they coexist in the same locations. The clonal evolution hypothesis explains better the cluster heterogeneity because each location in the tumor experience different microenvironment, thus develop different clusters of cells with adaptive cell phenotypes to the Darwinian selection. This study has a limited scope of presenting a new technique using a small sample size. This technique also requires a longer experimental and analysis time than only IHC or FISH tests. However, it can be potentially used in clinical context for additional assessments of equivocal or heterogeneous cases. In the future, a clinical study having a substantial number of cases and having a high number of ITH cases need to be performed to validate this technique and establish better the link between true ITH and cancer prognostics.

## Materials and methods

This section describes a brief summary of the materials and method. Detailed experimental protocols and mathematical background of this study can be found in the Supporting Information.

### Tissue selection

All cancer tissues were retrieved from the Institute of Pathology of Lausanne according to the ethical convention BB514/2012 established with the Ethical Commission of Clinical Research of the state of Vaud (Switzerland). All breast cancer patients did not oppose the use of their tissues for research purposes. Twenty formalin-fixed paraffin-embedded (FFPE) tissue samples were either primary breast cancers (16 cases) or metastatic breast cancer tissues in bone (2 cases) or stomach (2 cases). Thirteen selected cases were previously classified as equivocal by both IHC and FISH by a pathologist. Three cases were negative by FISH and equivocal/negative by IHC. Four other cases were HER2-positive by both FISH and IHC. The engineers (H.T.N. and D.M.) who performed the automatic analysis did not know the heterogeneity status of the samples. All tissues were assessed by a pathologist to determine the area of invasive ductal carcinoma using a Hematoxylin & Eosin stained section. An adjacent section, 4 µm in thickness, was processed for IF/FISH staining protocols.

### IF and FISH staining materials and protocols

The microfluidic staining was performed using rabbit anti-human c-erbB-2 oncoprotein primary antibody (HER2) and mouse anti-human CK (code A0485 and M3515, Agilent Technology, Basel, Switzerland) primary antibody mix and AF 594 and AF647 (code: A-11037 and A-21236, Thermo Fisher Scientific, Reinach, Switzerland) secondary antibody cocktail. FISH staining protocol is described in section S2 of the SI.

### Image processing for automatic FISH scoring

While the CEP17 signals are distinct and easily scored using local maximum detection, some small HER2 signals could overlap and form indistinguishable clusters in the low-resolution 20×-objective images. To estimate the number of *HER2* loci in each cluster, local maxima of the HER2 signal within a nucleus (cluster positions) were identified. Areas proportional to the full-width-half-maximum of each local maximum, corresponding to the area around the maximum, were then assigned to each cluster. The cluster size and integrated intensity (sum of all signal pixels within a cluster) were measured. The number of HER2 signals inside each HER2 cluster was estimated using a comparison between the number of red dots counted using a high-magnification (63×) objective and the integrated intensity of the HER2 cluster normalized to that of one HER2 dot signal.

### Cell selection and characterization

A selected nucleus had a size in the range of 50–1000 µm^2^ and contained at least one HER2 and one CEP17 signal. After this step, epithelial non-cancer cells are discarded. All non-epithelial cells having weak CK staining (CK signal less than Mean - 0.5 × standard deviation) are also discarded. Furthermore, epithelial non-invasive (*in situ*) cancer cells can be detected along with the invasive cancer cells, we manually select the regions where there are only invasive cells. For all remaining cells (epithelial invasive cancer cells), we computed the mean HER2/CK ratio, the number of *HER2* loci and *HER2*/CEP17 ratio per cell and their corresponding standard deviations. This mean cell *HER2*/CEP17 ratio correlates nearly-perfectly to the overall *HER2*/CEP17 ratio, which is the ratio between the sum of all *HER2* loci and CEP17 for all cells in the tissue (see section S4 of the SI). The threshold for positivity of cell *HER2*/CEP17 ratio is thus calculated from the overall *HER2*/CEP17 ratio threshold 2, giving 2.4.

### Analysis of local indication of spatial association (LISA)

The LISA method (detailed in section S5 of the SI) can quantitatively discriminate cluster and mosaic ITH by comparing the variable of interest for each cell with that of its neighbors. For ITH characterization, the two parameters used are the HER2/CK ratio and *HER2* loci/cell number, calculated for each cell and its neighborhood, in comparison to a high and low threshold, *T*_*h*_ and *T*_*l*_. For the HER2/CK ratio, *T*_*h*_ is equal to *T*_*l*_ (single threshold) and defined as the lower boundary of the confidence interval of a t-test on SKBR3 cell lines’ HER2/CK (*vide supra*), giving *T*_*h*_ =*T*_*l*_= 0.25. Cells and neighbors having HER2/CK ≥ 0.25 are classified as H, while they are classified as L if HER2/CK < 0.25. For the parameter *HER2* loci/cell, as this is a standard variable, we used the ASCO/CAP 2013 guideline to determine *T*_*h*_ and *T*_*l*_: *T*_*h*_ =6 and *T*_*l*_ =4 [1]. A cell having a *HER2* copy number > 6 is unambiguously positive and, when this number < 4, it is unambiguously negative. If the cell’s *HER2* copy number is between 4 and 6 or equal to 4 or 6, it is non-classified (NC) or equivocal. Neighborhoods are classified as High (respectively Low) if the mean of all neighborhood’s values is higher (respectively lower) than 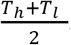.

### Statistical modeling of the *HER2*-loci-per-cell number in a homogeneous cell population

Mosaic heterogeneity can be biased by truncation artifacts because a positive cell can also display a smaller number of *HER2* loci/cell depending on the position of the cut and the probability that the *HER2* signals are placed inside the cut. Therefore, we simulated the truncation artifacts of a homogeneous cell population in function of the number of *HER2* loci/cell to distinguish these artifacts from the real mosaic ITH. To achieve this aim, first, we calculate the volume *V* _*τ*_(*x*) of the cell section from the relative position of the cut and the thickness of the cut *τ* in the cell is detailed in section S6 of the SI. For each *V* _*τ*_(*x*), we can calculate the probability *p*(*x*) for one *HER2* locus to be placed inside the cell section. We demonstrate that the number of *HER2* loci/cell follows a binomial law of probability. From *p*(*x*) we calculate the probability mass function of the *HER2* loci/cell variable (see section S7 of the SI) used for generating a truncated-homogeneous-cell population (see section S8 of the SI).

### Code availability

The software is uploaded to Github server (www.github.com/huutuannguyen/Image-processing-IF-FISH and www.github.com/huutuannguyen/tumor-heterogeneity-analysis). Data of the ITH cases are available in the repository Figshare with the identifier data DOI https://doi.org/10.6084/m9.figshare.6241796

## Acknowledgments

We thank the European Union Ideas program for supporting this work (Grant number ERC-2012-AdG-320404).

## Author contributions

H.T.N. designed the experimental plan, created image processing software, performed tissue analysis and built the heterogeneity theoretical model. Both H.T.N. and D.M. contributed to experimentation and data analysis. B.B. contributed to tissue assessments and case selection. Every author participated in scientific discussions. M.A.M.G. supervised the project. H.T.N wrote the manuscript, D.M, B.B., L.d.L, and M.A.M.G. revised the manuscript.

## Conflict of interest

Professor Martin A.M. Gijs has a patent application related to the microfluidic immunostaining (Swiss Patent Application 00256/12) and is involved in the startup Lunaphore technologies SA developing this technology.

